# Cell type specific novel lincRNAs and circRNAs in the BLUEPRINT haematopoietic transcriptomes atlas

**DOI:** 10.1101/764613

**Authors:** Luigi Grassi, Osagie G. Izuogu, Natasha A.N. Jorge, Denis Seyres, Mariona Bustamante, Frances Burden, Samantha Farrow, Neda Farahi, Fergal J. Martin, Adam Frankish, Jonathan M. Mudge, Myrto Kostadima, Romina Petersen, John J. Lambourne, Sophia Rowlston, Enca Martin-Rendon, Laura Clarke, Kate Downes, Xavier Estivill, Paul Flicek, Joost H.A. Martens, Marie-Laure Yaspo, Hendrik G. Stunnenberg, Willem H. Ouwehand, Fabio Passetti, Ernest Turro, Mattia Frontini

**Author notes:** Joint first authors (Luigi Grassi, Osagie G. Izuogu).

## Abstract

Transcriptional profiling of hematopoietic cell subpopulations has helped characterize the developmental stages of the hematopoietic system and the molecular basis of malignant and non-malignant blood diseases for the past three decades. The introduction of high-throughput RNA sequencing has increased knowledge of the full repertoire of RNA molecules in hematopoietic cells of different types, without relying on prior gene annotation. Here, we introduce the analysis of the BLUEPRINT consortium gene expression data for mature hematopoietic cells, comprising 90 total RNA and 32 small RNA sequencing experiments, from 27 different cell types. We used these data to describe the transcriptional profile of each we used guided transcriptome assembly to extend the annotation of the transcribed genome, which led to the identification of hundreds of novel non-coding RNA genes, which display a high degree of cell type specificity. We also characterized the expression of circular RNAs and found that these are also highly cell type specific. This resource refines the active transcriptional landscape of mature hematopoietic cells, highlights abundant genes and transcriptional isoforms for each cell type, and provides valuable data and visualisation tools for the scientific community working on hematological development and diseases.

## Introduction

Knowledge of the transcriptional programs underpinning the diverse functions of hematopoietic cells is essential to understand how and when these functions are performed and to aid the identification of the underlying causes of hematological diseases. Thanks to its accessibility, blood is the tissue of choice for the implementation of novel technologies in primary samples. Indeed, several studies aiming to characterise gene expression profiles have been performed on increasingly purified primary hematopoietic populations in the post genome era^1–3^. These studies used expression arrays and thus required prior specification of the sequences to be interrogated. The probed sequences were often derived from the analysis of a very limited number of tissues and cell types^4^, despite the early discovery that transcription is widespread throughout the human genome^5^. The introduction of high-throughput nucleic acids sequencing technologies^6^ has improved the assembly of the human genome and the annotation of transcriptomes therein, and it has enabled a much more comprehensive analysis of gene expression using transcriptomic assembly approaches^7^. The BLUEPRINT consortium^8^ was established to characterize the epigenetic state, including the transcriptional profile, of the different hematopoietic cell types. Reference datasets for DNA methylation, histone modifications and gene expression were generated using state-of-the-art technologies from highly purified cells, in accordance with quality standards set by the International Human Epigenome Consortium^9^. RNA sequencing (RNA-seq) data from over 270 samples encompassing 55 cell types have been made publicly available (http://dcc.blueprint-epigenome.eu), a subset of which has been described in other studies ^10–12^. Here, we present the analysis of 90 total RNA samples from 27 mature cell types from both cord and adult peripheral blood, together with 32 small RNA samples from 8 mature cell types. We used a Bayesian differential expression analysis approach^13^ to determine changes in the expression levels of genes and transcripts at lineage commitment events and to identify cell type specific transcriptional signatures. We performed guided transcriptome reconstruction^14^ using total RNA-seq reads, identifying 645 multi exonic transcripts originating from 400 intergenic novel genes. The majority of the novel transcripts have low protein coding potential and high cell type specificity. Additionally, we identified 55,187 circular RNAs (circRNAs), which also displayed very high cell type specificity, highlighting the emerging role of non-coding transcripts in hematopoiesis. To facilitate the exploration and reuse of the data by the biomedical community, we also provide an internet-based interface that allows to plot the expression patterns of genes and transcripts and to download normalised expression data (https://blueprint.haem.cam.ac.uk/bloodatlas/).

## Results

### The complexity of the hematopoietic transcriptomes

We isolated 90 samples (**Table S1**) from 72 whole blood and cord blood donations, either by magnetic beads separation or flow activated cell sorting (FACS; see M&M). We generated a mean of 91M 75bp paired-end reads for all total ribo-depleted RNA samples, except for platelets (PLT), basophils (BAS) and eosinophils (EOS), which we sequenced by 150bp paired-end sequencing at a comparable depth (**Table S1**). We also generated a mean of 4.5M 50bp single-end reads per small RNA sample (**Table S2**). Principal component analysis (PCA) of the log expression estimates for both long and short RNAs show distinct clustering by cell type according to their ontology along the first two principal components, which explain approximately 40% of the variance (Fig. 1A, 1B, S1A and S1B). This correspondence is also obtained by hierarchical clustering of samples using Spearman’s rank correlation across the samples (Fig. S1C and S1D).

**Figure 1:**
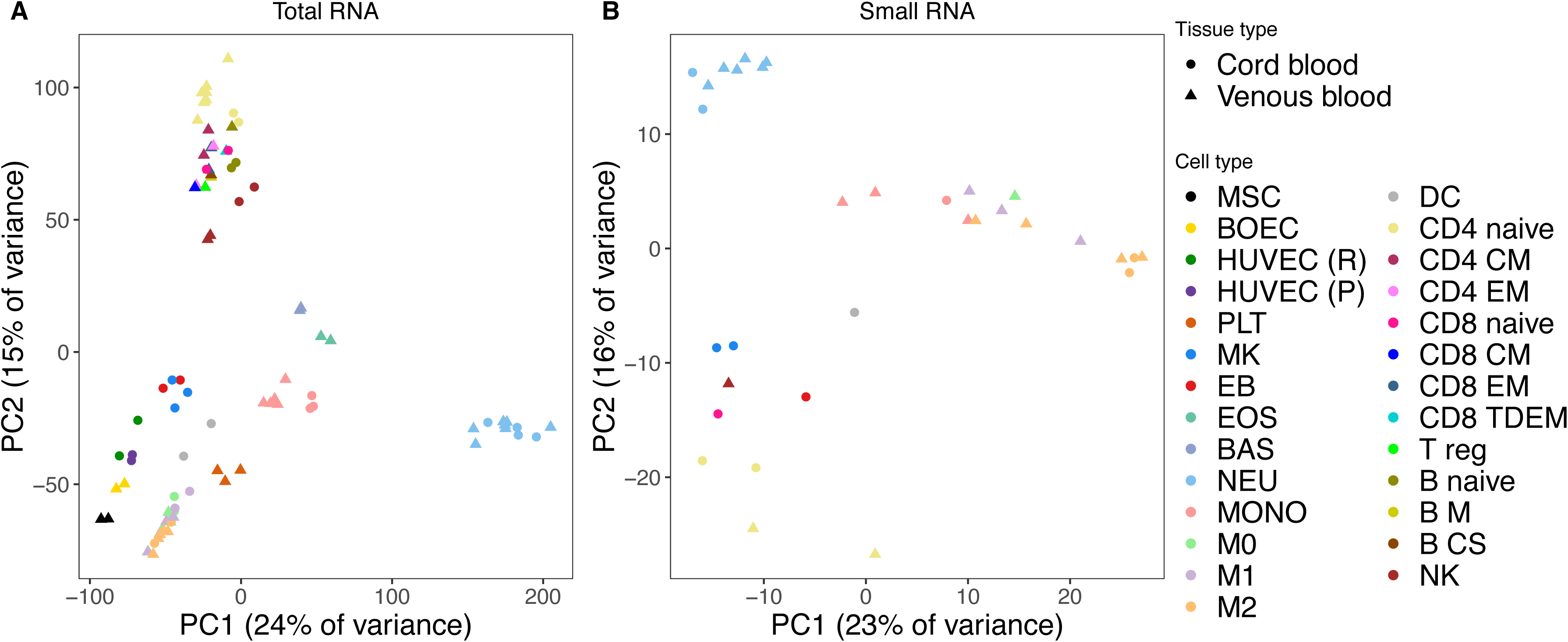
Principal component analysis of gene and miRNA expressions. **1A**: Scatterplot of the first (PC1) vs the second (PC2) principal component of the expression of genes with a log expression estimate greater than zero in at least one sample. **1B**: Scatterplot of PC1 vs PC2 of the expression of the miRNAs with unique read count >10 in at least one sample.

A fraction of the expressed genes typically dominate<u>s</u> the transcriptome of any given tissue or cell type in terms of amount of RNA molecules. The GTEx project^15^ has shown that whole blood, considered as a single tissue, has a very low gene expression complexity, with three hemoglobin genes contributing more than 60% of total reads^16^. We refined this analysis by studying transcriptome complexity in different cell types of blood. After excluding mitochondrial genes due to their considerable variation in steady-state expression across individuals^17^, the number of protein-coding genes accounting for 50% of expression ranged from only 14 in PLT to 600 in BAS. The number of protein-coding genes accounting for 75% of total expression ranged from 168 in PLT to 2,422 in resting human umbilical vein endothelial cells (HUVEC (R); Fig. 2A, Table S3, **Supplementary File 1**). For all cell types in this study, with the exception of PLT, the sets of genes yielding 75% of total reads showed gene ontology (GO) terms enrichment only for functional categories related to general biological processes, such as translation or transcription^18^. Thus, cellular integrity and basic cellular functions are supported at the transcriptional level even in mature cell types, some of which have short half-lives. In PLT, however, we found a GO terms enrichment for functional categories related to their core functions (i.e. hemostasis, wound healing, coagulation, platelet degranulation) while more general processes featured less prominently (**Table S4**). The small RNA landscape showed a very low complexity, with 50% of the reads in the 7 cell types originating from between 1 and 7 miRNAs (Fig. 2B, **Supplementary File 2**) and with fewer than 10 genes accounting for 75% of the RNA content in any cell type.

**Figure 2:**
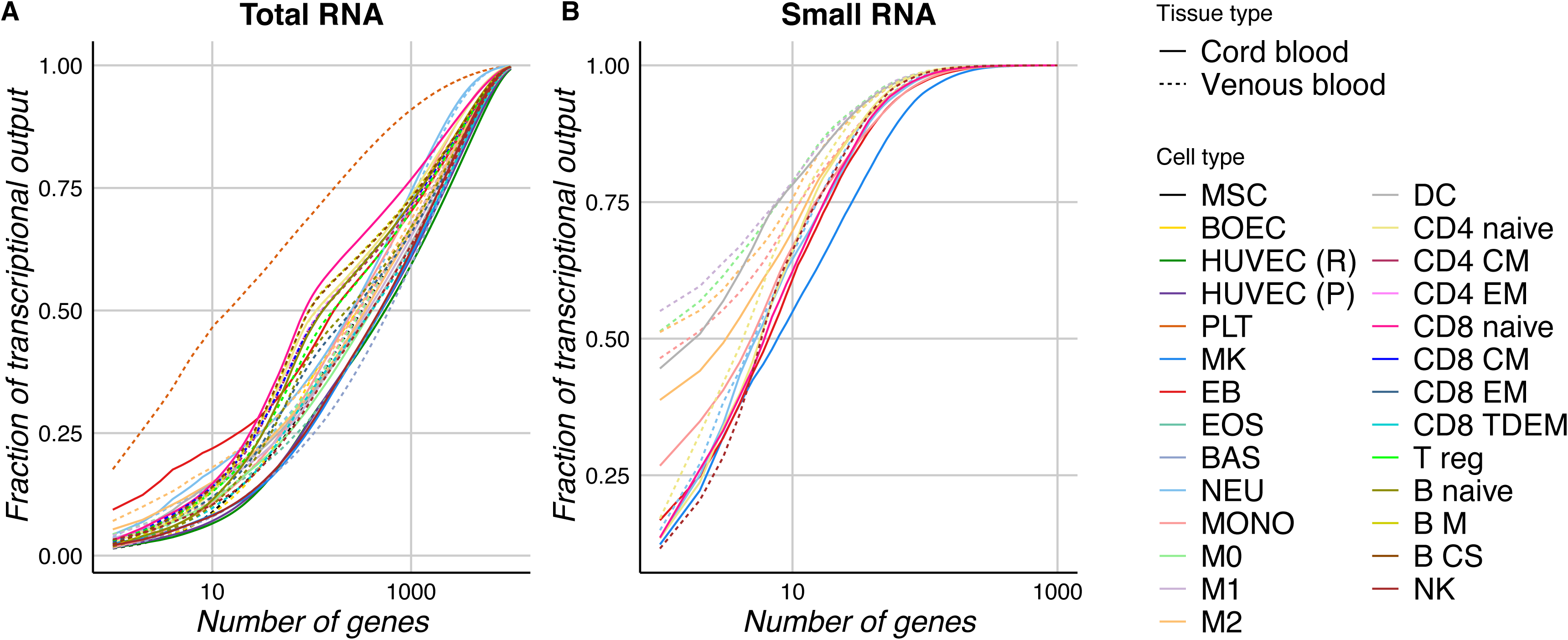
Complexity of genes and miRNA transcriptomes. **2A**: Cumulative distribution of the fraction of total transcription contributed by non-mitochondrial protein-coding genes when sorted from most to least expressed in each cell type. **2B**: Cumulative distribution of the fraction of total mature miRNA transcription contributed by mature miRNAs when sorted from most to least expressed in each cell type.

### Transcriptional signatures define hematopoietic cell functions

As the most highly transcribed genes in each cell type do not encode for the cell type’s specific functions, we reasoned that these functions must be encoded by genes which may not be highly expressed but which, nonetheless, have variable levels of expression across the hematopoietic tree. We identified heterogeneously expressed genes by comparing a statistical model having a global expression parameter across all cell types with one in which each cell type has its own expression parameter. Using this approach, we found 19,861 genes, representing 59.5% of all HGNC-annotated genes in Ensembl, that had a posterior probability of differential expression >0.8. The mean log expression across samples was >0 for over half of differentially expressed genes but only for 3.5% of non-differentially expressed genes, indicating that ubiquitously expressed housekeeping genes in haematopoiesis number in the few hundreds. The differentially expressed genes were then classified by the cell type with the greatest expression. To ensure that the signatures recapitulated cellular functions specific to the mature blood cells in this atlas, rather than functions of shared progenitors from which they originate, we subselected the 16,572 genes whose maximum log_e_ expression level was at least 0.1 (i.e. 10.5%) greater than that found in the cell type with the second greatest expression (**M&M**, **Table S5**). For example, VWF is tagged with the endothelial cells group label (ENDO) because its expression varies across cell types (posterior probability of the alternate model ~= 1), VWF is most highly expressed in ENDO (log_e_ expression level = 6.0), and the second highest expressed category (MK/PLT) has a log_e_ expression level of 2.2 (Fig. 3A). The number of genes assigned to each category ranged from 186 in CD8 T lymphocytes (CD8TC) to 3,502 in MK/PLT (Fig. 3B). Using these groups of genes, we found enrichment of GO terms reflecting the primary functions for all categories, except for BAS, macrophages M0 (M0) and monocytes (MONO), at a family-wise error rate < 5% (**Table S6**), as exemplified for the MK/PLT cluster and dendritic cells (DC) cluster in Fig. 3C.

**Figure 3:**
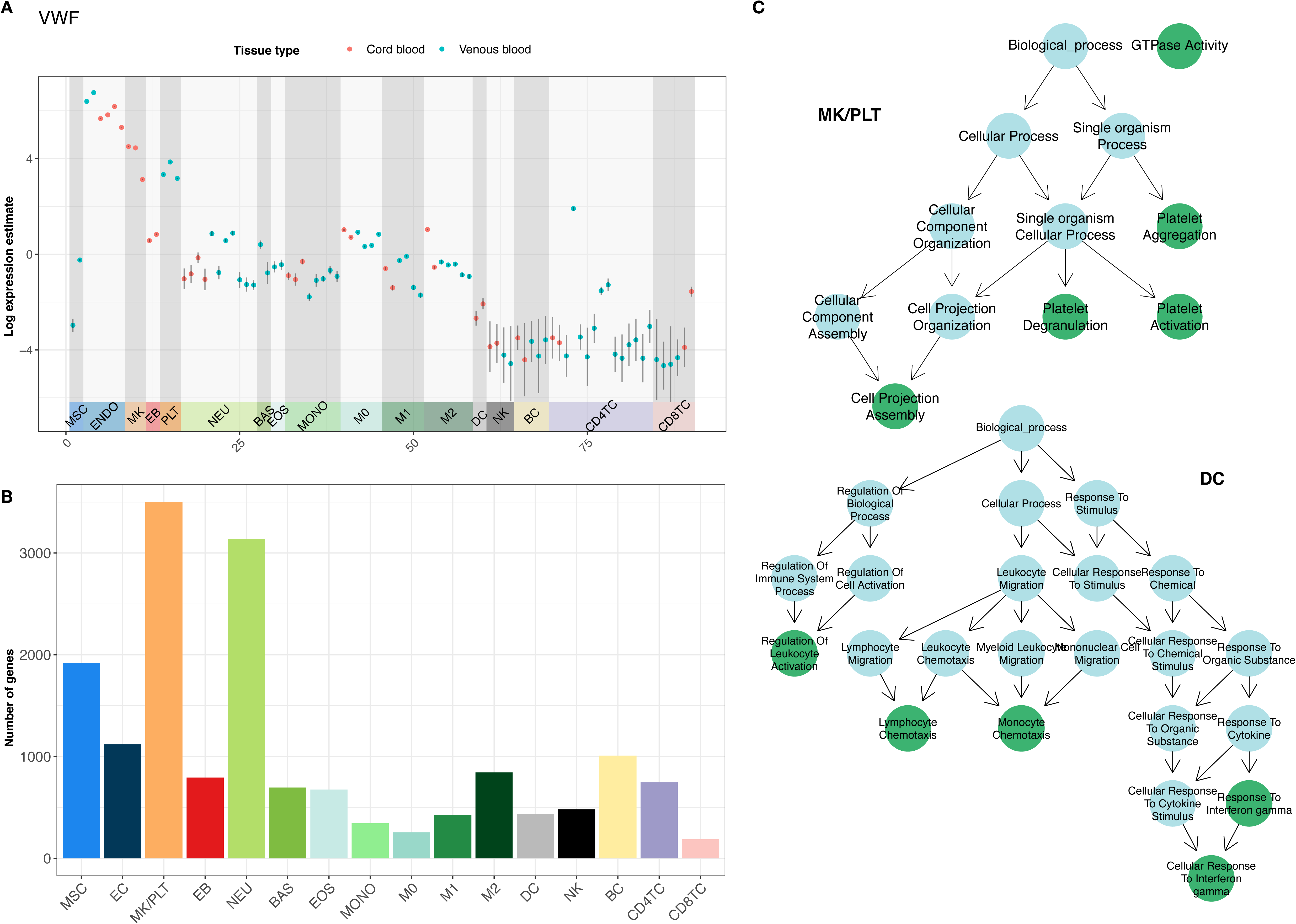
Cell type specific transcriptional signatures. **3A**: VWF expression estimates and posterior variances across samples. **3B**: The number of differentially expressed genes classified by cell type having the greatest expression, subject to a log fold change >0.1 compared to the cell type having the second greatest expression. **3C**: Graphical representations of the GO term enrichments for the MK/PLT and the DC groups in which the nodes represent terms, which are coloured green if they are enriched and light blue if they are ontological ancestors of enriched terms, and edges represent ontological relations.

### Differential expression of miRNAs

We applied the differential expression modelling described above to the short RNA data for four CD4TC, two MK, eight NEU, four MONO, three M1 and six M2 samples. We found 603 out of 2,588 miRBASE-annotated^19^ miRNAs to be differentially expressed with a posterior probability > 0.8, of which 573 exhibited a log_e_ fold change between the most highly expressed and the second most highly expressed cell type greater than 0.1 and were thus classified as cell type specific. The mean expression of miRNAs was strongly associated with their having at least one validated miRNA target amongst the 29,920 validated mRNA-miRNA interactions in the mirecords, mirtarbase and tarbase databases^20^ (P < 2 × 10^−16^, effect size = 0.16, logistic regression). For example, 46 of the 50 miRNAs (92%) having the highest mean expression over cell types had at least one validated target, while only 458/2508 (18.2%) of the remaining 2,508 miRNAs had a validated target. The cell type specific miRNAs with the greatest expression in their labelled cell type (**Table S7**) had been previously linked to relevant cellular functions. For example, hsa-miR-21-5p (the most highly expressed M1-specific miRNA) is involved in resolution of wound inflammation^21^ and macrophage polarization^22^; hsa-let-7g-5p, hsa-miR-26a-5p, hsa-miR-150-5p and hsa-miR-146b-5p (the most highly expressed CD4TC-specific miRNAs) are important modulators of CD4+ T-cells^23,24^; and hsa-miR-126-3p (the most highly expressed MK-specific miRNA) plays a role in MK/PLT biogenesis^25,26^. Using existing databases of miRNA-mRNA interactions, however, we did not find a correlation between expression of miRNAs and expression of their targets, which is consistent with miRNAs being only one of a diverse set of molecular players in transcriptional regulation of haematopoietic cells and in agreement with the results of other studies showing that miRNAs induce translational repression without mRNA destabilisation^27^.

### *De novo* transcriptome assembly identifies new genes and gene isoforms

The pervasive transcription of different types of non-coding RNAs (ncRNAs) is one of the most recent discoveries in the RNA biology of mammalian genomes^28^. Among ncRNAs, long ncRNAs (lncRNAs) form a heterogeneous class with crucial roles in the control of gene expression, both during developmental and during differentiation processes^29^. The number of lncRNA species is higher in the genome of developmentally complex organisms, hinting at the importance of RNA-based control mechanisms in the evolution of multicellular organisms^30^. Several studies have demonstrated that almost two-thirds of the genome is pervasively transcribed^31^. We used the BLUEPRINT gene expression dataset to identify novel genes and novel isoforms within known genes with respect to the reference transcriptome (Ensembl 75^32^). We constructed sample-specific transcriptomes using read alignments to the reference genome^33^ and merged them into a consensus transcriptome. The consensus transcriptome contained 645 multi-exonic transcripts from 400 novel genes, defined as genes that did not overlap any of the transcripts present in Ensembl 75, GENCODE 19^34^ or RefSeq^35^ (**Supplementary File 3**). We found that using the expression values of the 368 novel genes having a log expression >0 in at least one sample it was possible to cluster the different samples by cell type (Fig. 4A), suggesting these novel genes play a role in the determination of cellular identity.

**Figure 4:**
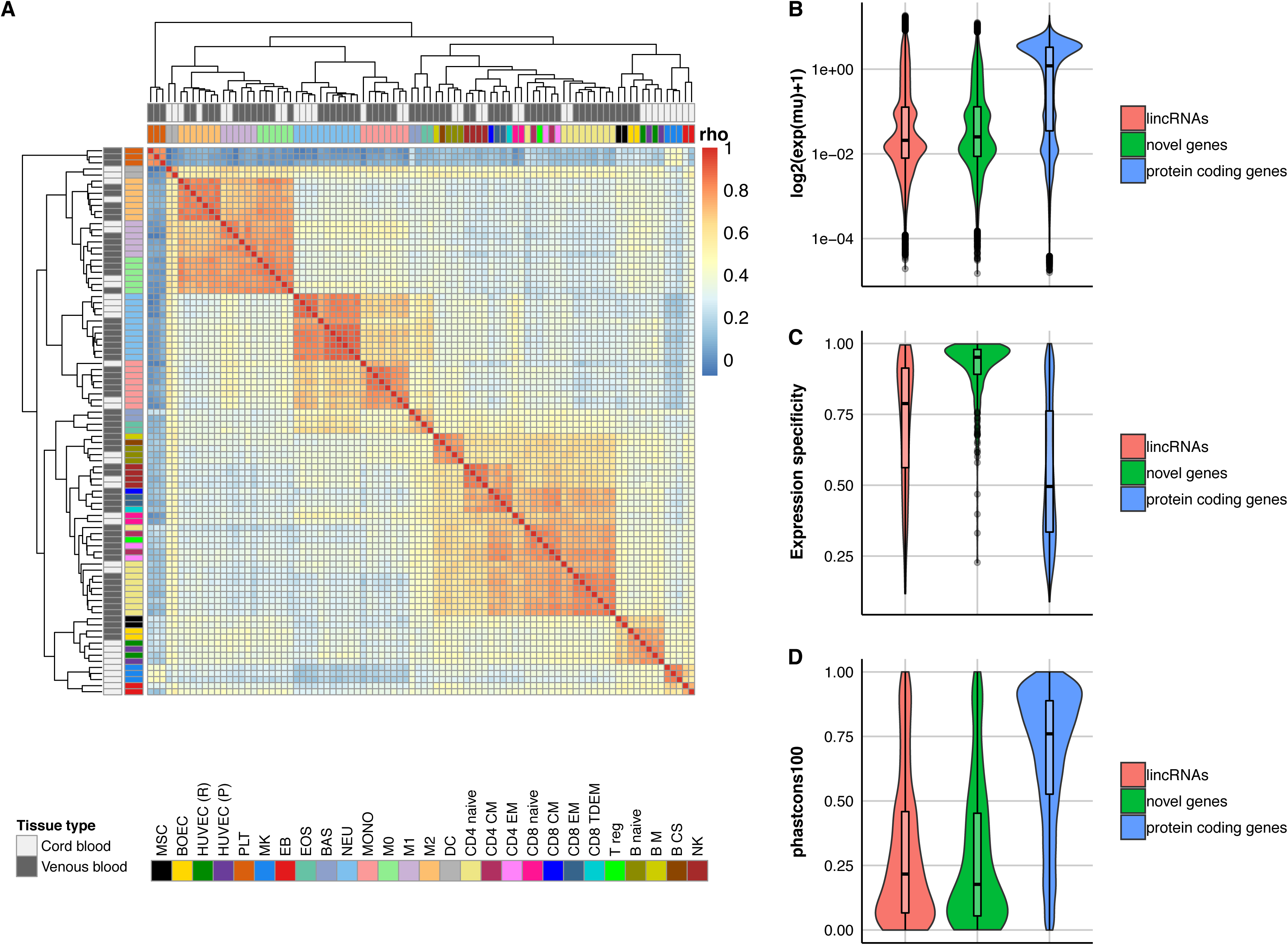
Properties of the identified novel genes. **4A**: Heatmap of the Spearman’s rank correlation (rho) matrix calculated by using the log2(FPKM+1) values of the 368 novel genes, expressed (FPKM>1) in at least one sample. Dendrogram has been drawn by using complete-linkage clustering based on distances calculated as one minus the correlation coefficient. **4B**: Expression distributions of the novel genes and the ones annotated in Ensembl 75 with biotype protein coding or lncRNAs. **4C**: Expression specificity (Tau) distributions of the novel genes and the ones annotated in Ensembl 75 with biotype protein coding or lncRNAs. **4D**: Sequence conservation (UCSC phastCons 100) distributions of the novel genes and the ones annotated in Ensembl 75 with biotype protein coding or lncRNAs. PhastCons have been obtained from multiple alignments of human (hg19) sequences with other 99 vertebrate species.

The vast majority (555 out of 645) of novel multi-exonic transcripts had a coding potential^36^ below 0.364, therefore classifying them as non-coding, whilst the remaining 90 transcripts were classified by CPAT as potentially coding. Additionally, to the CPAT score we also employed other discriminating features, such as the presence of low complexity regions, to separate coding from non-coding genes. Open reading frames (ORFs) annotated in GENCODE have indeed minimal overlap with transposon-associated regions and other repetitive or low complexity regions (~2 % of all nucleotide positions)^37^. To further investigate the coding potentials of this set of novel transcripts, we determined that the percentage of transcripts overlapping repeat elements in the non-coding and potentially coding categories is not significantly different (Fig. S2A), and that non-coding and potentially coding transcripts did not show differences in the portion of each transcript overlapping repeats regions (Fig. S2B) nor in the localization of the overlap with repeat regions (Fig. S2C). These findings indicate that amongst the novel genes, even those with a higher coding potential display features that are more similar to non-coding transcripts rather than protein coding ones, for this reason we chose not to separate the two groups. Furthermore, the distribution of the expression level of the novel genes is lower than that of known protein coding genes (Ensembl 75) and it is similar to that of annotated lncRNAs (Fig. 4B). Novel genes also have higher tissue specificity than known lncRNAs and protein coding genes annotated in Ensembl 75 (Fig. 4C). These two properties contribute to explain their novelty: novel genes are expressed only in a very limited number of cell types and at low level, albeit consistently across biological replicates. Therefore, their identification has been made possible only upon the reconstruction of cell type specific transcriptomes. Supporting their prevalent non-coding nature is also the poor conservation of the exonic sequences across vertebrates, again resembling that of annotated lincRNAs, rather than that of protein coding genes (Fig. 4D). The genomic coordinates of these novel genes are available as a supplementary gtf file (**Supplementary file 3**).

### Circular RNA in mature hematopoietic cells

Circular RNAs (circRNAs) are single stranded RNA molecules whose ends are covalently joined via a back-splice mechanism. Most circRNAs have unknown function but some circRNAs are known to regulate transcription^38^ or act as miRNA sponges^39–41^. Peripheral blood contains thousands of circRNAs expressed at higher levels than their corresponding linear mRNAs^42^. We determine the abundance of circRNAs in the total RNA-seq data using five methods^40,43–46^; requiring that each identified backsplice event is detected by at least three methods to mitigate aligner-specific biases and exclude predictions that overlap known segmental duplications^47^ in the genome, multiple genes or Ensembl 75-annotated readthrough transcripts. We obtained a final list of 91,866 circRNAs, 55,187 of which were observed in multiple samples (**Supplementary Table 8**). The vast majority (81.64%) of back-splice events we identified were exonic and utilized annotated canonical splice sites (Fig. 5A), as expected from previous reports^40,48^. Many (44%) of the circRNAs matched structures in circBase^49^ exactly, and a further 30% overlapped structures in circBase. In comparison to other RNA species, circRNAs have low abundance, but they can accumulate inside the cell as a result of their resistance to exonuclease activity^50^. To investigate the expression patterns of circRNAs in the different hematopoietic cells, we performed pairwise correlation analysis and hierarchical clustering of Spearman’s correlation coefficients using only counts from circRNAs observed in multiple samples. These analyses distinctly grouped samples by cell types and lineage, to show tissue-specific expression of circRNAs (Fig. 5B). Next, we compared circRNA abundance with the expression of the linear RNAs originating from same genes, using as measure abundance ratios (AR), calculated by dividing the back-splice read counts from each locus with the canonical junction counts. We found mean ARs over replicates within cell types ranging from 1.02% in HUVEC (R) to 12.45% in PLT (Fig. S3A and **Supplementary Table 9**); the latter due to the absence of steady-state transcription, in the anucleated PLT, and to the differential decay of circRNA relative to linear molecules^51^. We also observed that in 74.53% of genes producing circRNAs (n = 9,277), expression profiles of backsplice and canonical junctions from same loci are positively correlated (median rho: 0.13; interquartile range (IQR): 0.29) across cell types. Furthermore, over a third of these (38.04%) exhibit significant correlation between expressions of circRNAs and linear molecules. For this subset, the median expression of backsplice and canonical junctions are significantly higher (p-value < 2.2e-16 and p-value = 5.757e-13 respectively, Wilcoxon rank sum test), relative to other circRNA genes (Fig. 5C). Without ruling out the possibility that the small difference in median canonical junction expression is influenced by junctions internal to circRNAs from the same loci, it is conceivable that small changes in the transcriptional output of some genes results in higher observable circRNA expression due to their accumulation. Finally, to identify differentially expressed circRNA, we performed pairwise comparisons of their abundance and identified 984 distinct circRNAs (<2%), originating from 751 genes (protein-coding: 731; non-coding: 20) as differentially expressed. The maximum number of differentially expressed circRNAs observed from pairwise comparisons is 314 (median: 24, IQR: 48; Fig. S3B and **Supplementary File 4**). Moreover, the expression patterns of differentially expressed circRNAs cluster samples by cell type (Fig. 5D). Although several mechanisms of action have been discovered for non-coding RNAs, only a handful of circRNAs have been experimentally verified as functional^38,40^ and their functions are distinct from those of their host genes, negating direct functional inferences from GO analysis.

**Figure 5:**
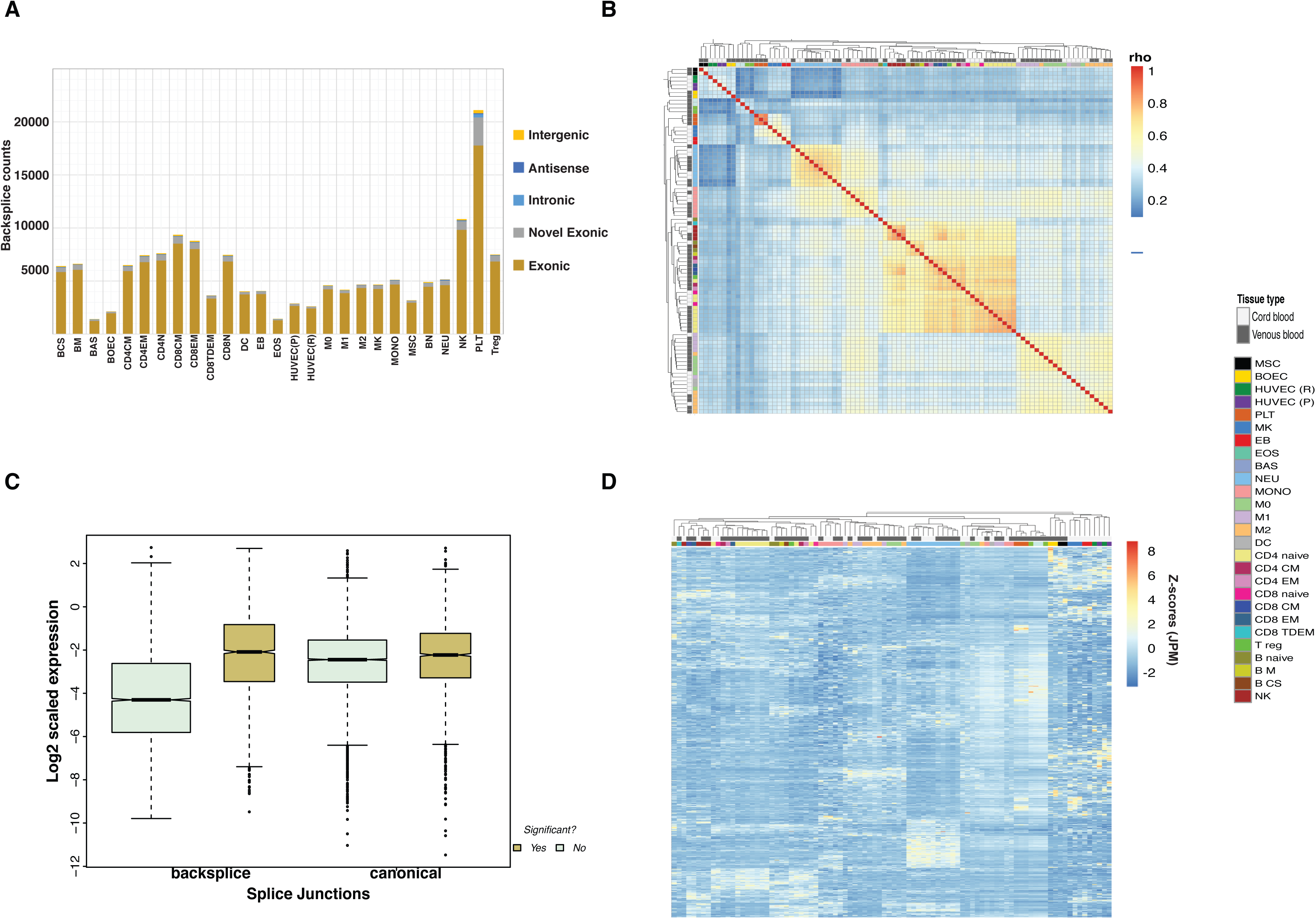
CircRNA expression in blood cells. **5A**: Bar plot showing distributions of circRNAs identified from all samples, grouped by cell types. Each bar is colour-coded to indicate number of identified circRNAs originating from different genomic regions. **5B**: Heatmap of the Spearman’s rank correlation (rho) using back splice junction counts from each sample. Lowly expressed circRNAs (with < 20 reads from all samples) were excluded. **5C**: Boxplots showing distributions of splice junction expression in circRNA producing genes. Boxes are colour-coded to show splice junction expression distributions in genes with correlation between circRNA and linear RNA abundance. **5D**: Heat map showing expression of all differentially expressed circRNAs (n = 987) identified from pairwise comparisons between cell types.

## Discussion

Here we explored 90 transcriptomes, from mature hematopoietic cells produced by the BLUEPRINT consortium, with the aim to determine which genes allow each of the 27 cell types achieves their diversity (Fig. 1) and their unique functional role in the hematopoietic system. We have shown that, at best, 2422 genes (ranging from 168 to 2422), out of the ~10,000 considered expressed at >=1 FPKM or more, form 75% of each transcriptome and that these are enriched in genes encoding for proteins involved in basic cellular functions rather than in those required to specify the different functional phenotypes/identities, the only exception being platelets, which have a much simpler transcriptome, 75% of which is occupied by 168 genes encoding for their core functions (Fig. 2). For the remaining cell types functional identity is achieved by the establishment of expression patterns, composed of uniquely expressed genes and of genes whose expression level is differing in the various samples (Fig. 3). These were identified using a differential expression analysis deploying a Bayesian statistical model (M&M). We conclude that each hematopoietic cell type performs its functions by expressing a unique combination of genes, partially overlapping with other cell types, and that basic cellular functions are up kept even in cell types with very limited half-life. Next, we leveraged on RNA-seq annotation agnostic nature to use genome alignments to reconstruct the transcriptome of each cell type and to identify, with a very conservative approach, at least 400 novel genes. These display properties such as low expression and high tissue specificity, that are highly reminiscent of those of lncRNAs^52^ (Fig. 4). The nature of the data (ribo-depletion) allowed also to greatly expand the catalogue of circRNAs identified in blood, as well as, to determine that these ncRNAs display high levels of cell type specificity (Fig. 5). Our findings reinforce the notion that lncRNAs and circRNAs may have roles in determining cell fate and functions in hematopoiesis, through mechanisms that are yet to be investigated.

Finally, our website https://blueprint.haem.cam.ac.uk/bloodatlas/ provides an interface for exploring expression levels at the gene and transcript levels generating graphical representations and downloading expression values.

## Acknowledgments and funding

The authors would like to the participation of National Institute of Health Research (NIHR) Cambridge BioResource volunteers and thank the NIHR Cambridge BioResource staff for their support. The work was funded by a grant from the European Commission 7th Framework Program (FP7/2007–2013, grant 282510, BLUEPRINT) to XE, PF, JHAM, MY, HGS and WHO. WHO is an NIHR senior investigator and receives funding from Bristol-Myers Squibb, the British Heart Foundation, the Medical Research Council and the National Institute for Health Research (NIHR). OGI, FJM, AF, JMM, LC and PF are funded by the Wellcome Trust (WT108749/Z/15/Z) with additional funding for specific project components such as GENCODE from the National Human Genome Research Institute of the National Institutes of Health (2U41HG007234), accordingly the content of this manuscript is solely the responsibility of the authors and does not necessarily represent the official views of the National Institutes of Health. KD is a HSST trainee supported by NHS Health Education England. NF is funded by the National Institute for Health Research (NIHR) Cambridge Biomedical Research Centre. FP is supported by the Fundação Carlos Chagas Filho de Amparo à Pesquisado Estado do Rio de Janeiro (FAPERJ; E-26/203.229/2016). NANJ is a recipient of a scholarship from the Coordenação de Aperfeiçoamento de Pessoal de Nível Superior - Brasil (CAPES; Finance Code 001). DS work has been supported in part by an Isaac Newton fellowship to MF. MF is supported by the British Heart Foundation (FS/18/53/33863).

## Materials & Methods

### Cell isolation

Samples were obtained from NHS Blood and Transplant donors and from cord blood donations at Cambridge University Hospitals, after informed consent (REC East of England 12/EE/0040). See supplementary material.

### RNA extraction

RNA was extracted from TRIzol according to manufacturer’s instructions, quantified using a Qubit RNA HS kit (Thermofisher) and quality controlled by Bioanalyzer (Agilent).

### Library construction

Libraries were prepared with TruSeq Stranded Total RNA Kit with Ribo-Zero Gold (Illumina) except for platelet, eosinophil and basophils which were prepared with Kapa stranded RNA-seq kit with riboerase (Roche).

### miRNA extraction

RNA was extracted with miRNeasy Mini Kit (Qiagen) and libraries prepared with NEBNext® Multiplex Small RNA Library kit (New England Biolabs).

### Expression analysis

Read were trimmed with Trim Galore (v0.3.7; parameters “-q 15 -s 3 --length 30 -e 0.05”) and aligned to Ensembl v75^7^ human transcriptome with Bowtie^53^ (1.0.1; parameters “-a --best --strata -S -m 100 -X 500 --chunkmbs 256 --nofw --fr”). MMSEQ^13,54^ (v1.0.10; default parameters) was used to quantify and normalise expression.

### Guided transcriptome assembly

STAR (v2.4.1c) with parameters “--runThreadN 8 --outStd SAM --outSAMtype BAM Unsorted --outSAMstrandField intronMotif” was used to align trimmed reads to Ensembl v75 human genome. The bam files sorted by coordinate and indexed by using samtools (v 1.3.1)^55^ were used for the guided transcriptome assembly with stringtie (v 1.3.4)^14^ with the parameters “-p 8 --rf -G Ensembl_75.gtf -v -l BPSTRG” and Ensembl v75 gtf as reference. Stringtie was used to merge individual transcriptomes in the master transcriptome. Gffcompare^56^ was used to compare the master transcriptome to the reference (Ensembl 75). Intergenic transcripts were further compared with gencode (v19)^34^ and ucsc (v hg19)^57^ transcriptomes by using the R bioconductor GenomicRanges package^58^, in order to exclude any overlap with those. Protein coding potential of the novel intergenic multi-exonic transcripts was assessed by using CPAT (v 1.2.4)^36^.

### CircRNA identification and expression profiling

A detailed description of computational methods for circRNA identification, expression profiling and comparisons is in the supplementary materials.

---

**Figure S1:**
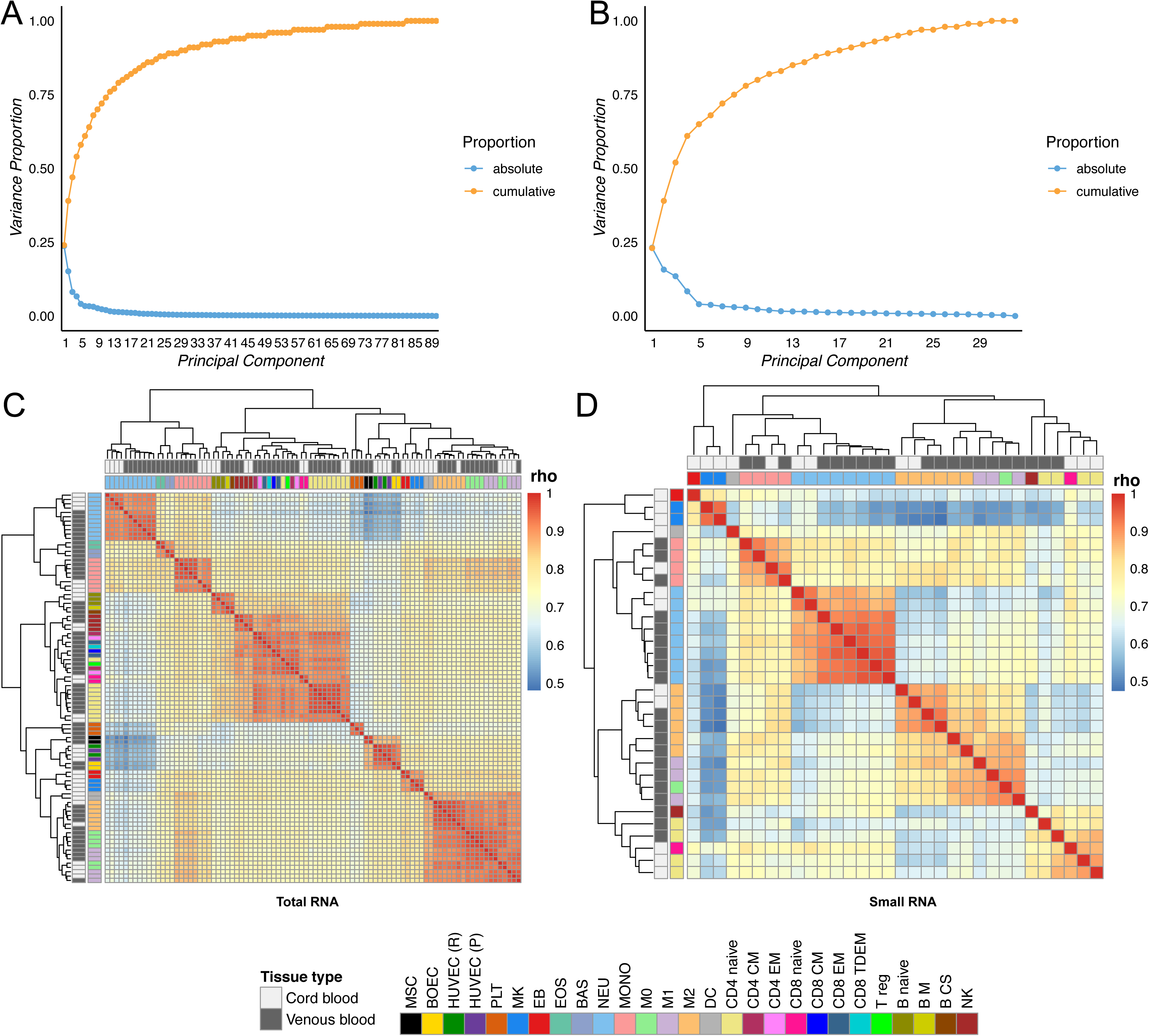
Gene and miRNA expressions PCA and correlation clustering. **1A**: Cumulative variance plot for each principal component. Genes with a log expression estimate greater than zero in at least one sample have been included.**1B**: Cumulative variance plot for each principal component. miRNAs with unique read count >10 in at least one sample have been included. **1C**: Heat map of the Spearman rank correlation coefficient (rho) between samples. Genes with a log expression estimate greater than zero in at least one sample have been used to calculate the Spearman rank correlation coefficient. Rows and columns order reflects the result of the complete linkage clustering made by using 1-rho as distance. **1D**: Heat map of the Spearman rank correlation coefficient (rho) between samples. MiRNAs with unique read count >10 in at least one sample have been used to calculate the Spearman rank correlation coefficient. Rows and columns order reflects the result of a complete linkage clustering made by using 1-rho as distance.

**Figure S2:**
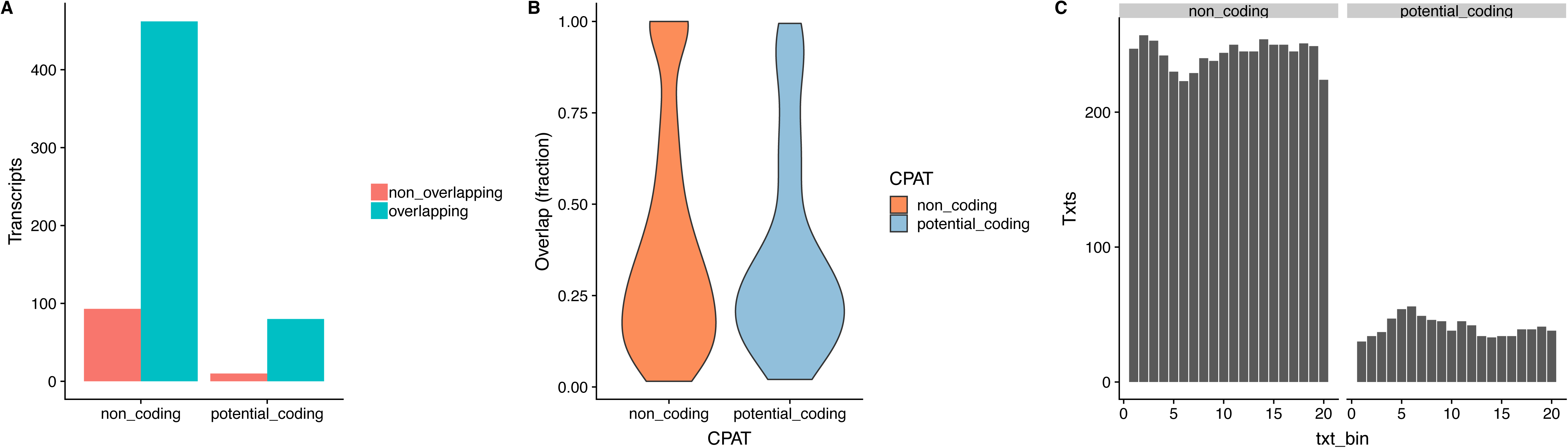
Repeat elements overlap of novel transcripts. **2A**: Non coding and potential coding transcript have non significant difference in the fraction of transcripts overlapping repeats (Fisher’s Exact Test p-value = 0.2147, odds ratio 0.6214023). **2B**: Non coding and potential coding transcripts have similar fraction of overlapping repeats, wilcoxon test (#W = 19316, p-value = 0.5184).

**Figure S3:**
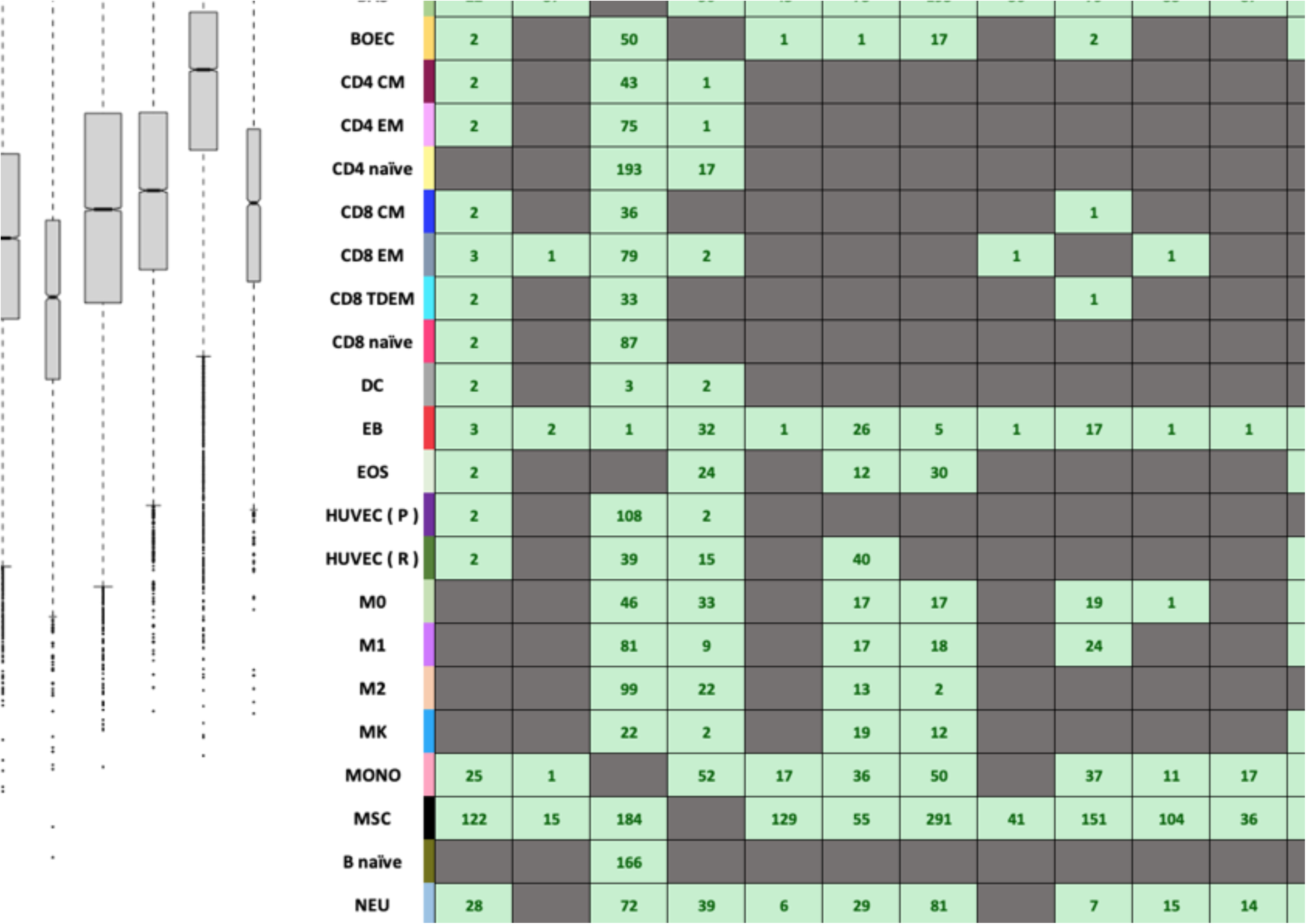
Differential circRNA abundance in blood cells. **3A:** Box and whisker plots showing distributions of circRNA abundance ratios in blood cells. Abundance ratios were derived by dividing back-splice junction counts with total splice reads from host genes. **3B**: Heatmap indicating numbers of differentially expressed circRNAs identified from pairwise comparisons of circRNA expression in blood cells.

## Extended methods

### Materials & Methods

#### Cell isolation

All samples were obtained from NHS Blood and Transplant blood donors and processed within 3 hours, and from cord blood donations at Rosie Hospital, Cambridge University Hospitals, in both cases after informed consent (ethical approval REC East of England 12/EE/0040). Detailed protocols, including antibodies panels, have been made available at http://www.blueprint-epigenome.eu/. Briefly neutrophils and monocytes were isolated from peripheral blood whole units (460 ml) of or from cord blood units. Peripheral blood mononuclear cells (PBMCs) were separated by gradient centrifugation (Percoll 1.078 g/ml) whilst neutrophils were isolated from the pellet, after red blood cell lysis, by CD16 positive selection (Miltenyi). PBMCs were further separated using a second gradient (Percoll 1.066 g/ml) to obtain a monocyte rich layer. Monocytes were further purified by CD16 depletion followed by CD14 positive selection (Miltenyi). For neutrophils and monocytes gene expression was tested also on Illumina HT12v4 arrays (accession E-MTAB-1573 at arrayexpress). The purification of macrophages M0, LPS activate macrophages M1, alternatively activated macrophages M2, endothelial cell precursors, erythroblasts, megakaryocyte, naive B lymphocytes, naive CD4 lymphocytes, naive CD8 lymphocytes used in this study has been extensively described{25258084}{28703137}. Regulatory CD4 lymphocytes (T regs), CD4 central memory lymphocytes (CM) and CD4 effector memory lymphocytes (EM) were isolated by flow activated cytometry (FACS) using the following surface markers combinations: T regs, CD3+ CD4+ CD25+ CD127low; CD4 CM, CD3+ CD4+ CD45RA-CD62L+; CD4 EM, CD3+ CD4+ CD45RA-CD62L-. CD8 central memory lymphocytes (CM), CD8 effector memory lymphocytes (EM) and CD8 terminally differentiated effector memory lymphocytes (TDEM) were isolated by FACS using the following surface markers combinations: CD8 CM, CD3+ CD8+ CD62L+ CD45RA-; CD8 EM, CD3+ CD8+ CD62L-CD45RA-; CD8 TDEM, CD3+ CD8+ CD62L-CD45RA+. B memory lymphocytes and B class switch lymphocytes were isolated by FACS using the following surface markers combinations: B memory, CD19+ CD27+ IgD+; B class switch, CD19+ CD27+ IgD-CD38dim. Natural Killer cells (NK) were isolated by FACS using the following surface markers: CD3-CD56dim CD16+. Eosinophils and basophils were isolated from a mixed leukocytes pellet obtained by sedimentation of whole blood 6% hydroxyethyl starch (Grifols, Cambridge, UK) for 30 minutes using Easysep (Stemcell Technologies) as previously described{20805156}. Monocyte derived dendritic cell were generated from cord blood CD34 depleted PBMC after a second Percoll (1.066 g/ml) to enrich for monocytes using a PromoCell dendritic cell isolation kit. Bone marrow derived mesenchymal stem cells isolation had been previously described{18557828}. Platelets were isolated from platelet rich plasma after leukocyte (CD45 positive) depletion as previously described{28703137}. All cell types purity was assessed by flow cytometry and/or morphological analysis after cytospin preparations were made and stained. The purified cells were resuspended in Trizol. Samples which did not meet predefined criteria (>95%) of cell purity were not sent for data generation.

#### RNA extraction

RNA was extracted from TRIzol according to manufacturer’s instructions, quantified using a Qubit RNA HS kit (Thermofisher) and quality controlled using a Bioanalyzer RNA pico kit (Agilent).

#### Library construction and sequencing

For all cell types with the exceptions of platelet, eosinophil and basophils libraries were prepared using a TruSeq Stranded Total RNA Kit with Ribo-Zero Gold (Illumina) using 200ng of RNA as input. Platelet, eosinophil and basophils samples were prepared with the Kapa stranded RNA-seq kit with riboerase (Roche) according to the manufacturer’s instructions.

#### miRNA extraction

RNA was extracted using the miRNeasy Mini Kit (Qiagen) from cell pellets with an RNA Integrity Numbers (RINs) from 7.3 to 10 as assessed with an RNA 6000 Nano kit on a 2100 Bioanalyzer (Agilent). Small RNA libraries were prepared using the NEBNext® Multiplex Small RNA Library Prep Set for Illumina (New England Biolabs) and the LongAmp Taq 2x Master Mix. Size selection was performed with 6% polyacrylamide gels, and library quality was verified on a 2100 Bioanalyzer (Agilent). Equimolar (2 nM) amounts of each library, as verified with Picogreen® dsDNA Quantification Reagent (Promega), were pooled and sequenced on an Illumina HiSeq 2000 using 50 bp single end reads.

#### Expression analysis

Trim Galore (v0.3.7) (http://www.bioinformatics.babraham.ac.uk/projects/trim_galore/) with parameters “-q 15 -s 3 --length 30 -e 0.05” was used to trim PCR and sequencing adapters. Trimmed reads were aligned to the Ensembl v75 {25352552} human transcriptome with Bowtie 1.0.1 {19261174} using the parameters “-a --best --strata -S -m 100 -X 500 --chunkmbs 256 --nofw --fr”. MMSEQ (v1.0.10) {24281695}{21310039} was used with default parameters to quantify and normalise gene expression.

#### Guided transcriptome assembly

STAR (v2.4.1c) with parameters “--runThreadN 8 --outStd SAM --outSAMtype BAM Unsorted --outSAMstrandField intronMotif” was used to align trimmed reads to the Ensembl v75 (Cunningham et al., 2015) human genome. The bam files sorted by coordinate and indexed by using samtools (v 1.3.1) {19505943} have been used for the guided transcriptome assembly by using stringtie (v 1.3.4) {25690850} with the parameters “-p 8 --rf -G Ensembl_75.gtf -v -l BPSTRG” and the Ensembl v75 (Cunningham et al., 2015) gtf as reference transcriptome. Stringtie has also been used to merge the transcriptomes of each individual sample in one single master transcriptome. Gffcompare{22383036} has been used to compare the master transcriptome to the reference transcriptome (Ensembl 75). Intergenic transcripts have been further compared with gencode (v19){22955987} and ucsc (v hg19) {25428374} transcriptomes by using the R bioconductor GenomicRanges package {23950696}, in order to exclude any overlap with other annotated transcriptomes. The protein coding potential of the novel intergenic multiexonic transcripts has been assessed by using CPAT (v 1.2.4){23335781} using default parameters and human models provided by the program.

#### CircRNA identification and expression profiling

##### Identification and comparisons

Back-splice junctions were identified using CIRI{25583365}, CIRCexplorer{27365365}, find_circ{23446348}, circRNA_finder{25544350} and PTESFinder{26758031} (parameters: JSpan=10, PID=0.85, segment_size=65), mapping against the human genome (GRCh37). Candidate circRNA junctions were selected if reported by at least 3 methods and do not overlap segmental duplications. Genomic positions of back-splice junctions were compared to previously identified junctions in circbase.org{25234927} (obtained 05/2018), annotated splice sites in Ensembl 75{25352552} and known segmental duplications{11381028} in the genome. Back-splice junctions overlapping multiple genes, readthrough transcripts and duplicons were excluded from downstream analyses.

##### Classification

Identified circRNAs were classified into 5 groups based on their genomic location relative to Ensembl 75 annotations and overlap with known splice sites. *exonic_known:* Splice junction corresponds to known splice sites; *exonic_novel:* back-splice overlaps at least one annotated exon and utilizes only one known splice site; *intronic:* circRNA is internal to annotated intron; *intergenic:* back-splice junctions do not overlap annotated exons/introns and *antisense:* circRNAs overlap antisense to annotated exons/introns.

##### Expression estimates

Raw counts reported by PTESFinder were normalized by dividing with the total splice reads from each sample and multiplied by 1E6 to derive Junctions Per Million (JPMs). Abundance ratios were derived by dividing total back-splice reads with total spliced reads from each circRNA producing gene. Across all samples, z-scores of mean circRNA and canonical junction expression were compared to assess correlation. Statistical analysis of circRNA expression was performed using DESeq2{25516281}.

